# An extended Cks1-interacting region in Far1 modulates Cdk1 inhibition and substrate phosphorylation

**DOI:** 10.64898/2026.11.13.766371

**Authors:** Artemi Maljavin, Juli Mukhadze, Viacheslav Kiselev, Ervin Valk, Marko Lõoke, Mihkel Örd, Ilona Faustova

## Abstract

Cyclin-dependent kinases (CDKs) use multiple docking interactions to recognise their substrates and inhibitors, but how sequences between canonical interaction motifs contribute to these processes remains poorly understood. Here, we identify an auxiliary interaction element in the budding yeast CDK inhibitor Far1 and investigate its contribution to kinase inhibition and multisite phosphorylation. Combining AlphaFold3 modelling with biochemical assays and mutational analysis, we show that Far1 residues 310–319 (QLIRTADISC), located immediately downstream of the Cks1-binding phospho-threonine T306, contribute to inhibition of Clb5–Cdk1. In particular, the hydrophobic residues I312 and I317 are important for inhibitory potency, consistent with their predicted positions at the Cks1–Cdk1 interface. To investigate the functional consequences of this interaction in a substrate context, we introduced the Far1 sequence between two phosphorylation sites in a Sic1-derived substrate. This insertion increased the accumulation of doubly phosphorylated product by more than twofold, while reducing the overall phosphorylation rate. These findings suggest that sequences between canonical CDK recognition elements can provide auxiliary interactions that influence both kinase inhibition and substrate phosphorylation dynamics. Thus, the region between established docking motifs represents an additional determinant of CDK substrate processing and inhibitory activity.

## Introduction

Cyclin-dependent kinase (CDK) complexes phosphorylate hundreds of proteins in precise order to drive the cell cycle, and both the kinase and its regulation are conserved from yeast to humans (Morgan, 2006; Nurse and Bissett, 1981; Örd and Loog, 2019; Ubersax et al., 2003). CDK complex has three subunits: the catalytic kinase, a regulatory cyclin that mediates substrate protein recruitment, and the phosphoadaptor Cks1 (Bourne et al., 1996; Reynard et al., 2000). The kinase subunit recognises full S/T-P-x-K/R, minimal S/T-P consensus, and in some contexts non-proline directed phosphosites (Örd and Loog, 2019; Valk et al., 2023). Additional substrate specificity arises from docking interactions with the hydrophobic patch and other pockets on cyclins, and the phospho-threonine binding pocket of Cks1 (Bandyopadhyay et al., 2020; Hao et al., 2005; Loog and Morgan, 2005; McGrath et al., 2013; Ng et al., 2025; Örd et al., 2020, 2019). These pockets recognise distinct short linear motifs (SLiMs)in the intrinsically disordered regions (IDR) of CDK substrates and inhibitors.

In budding yeast, the four major cyclin pairs bind to four different motif classes: G1/S-phase Cln1/2 recognise LP motifs, S-phase Clb5/6 as well as human cyclins bind RxL motifs, G2-phase Clb3 recognises proline-rich PxxPxF motifs, and M-phase Clb2 binds LxF motifs (Bandyopadhyay et al., 2020; Bhaduri and Pryciak, 2011; Örd et al., 2020, 2019).Phosphorylation can also create docking sites, as Cks1 binds phospho-threonines to promote multisite phosphorylation (Kõivomägi et al., 2013; McGrath et al., 2013).

Cyclin-dependent kinase inhibitor proteins (CKIs) arrest the cell cycle by binding the complex directly using many of the same motifs as CDK substrates. In budding yeast, Sic1 blocks Clb5,6-Cdk1 at the G1/S transition (Kuhn et al., 1993; Schwob et al., 1994), the N-terminal domain of Cdc6 inhibits Clb2-Cdk1 during mitotic exit (Calzada et al., 2001; Elsasser et al., 1996), Far1 inhibits Cln1,2-Cdk1 and Clb5,6-Cdk1 downstream of the pheromone pathway (Gartner et al., 1998; Peter and Herskowitz, 1994), and Cip1 inhibits Cln-Cdk1 in response to stress (Chang et al., 2017; Ren et al., 2016). These four CKIs share no sequence homology and are largely intrinsically disordered. While we lack structural understanding of the inhibitory mechanism of yeast CKIs, more is known about the two families of human CKIs. INK4 proteins (p16, p15, p18 and p19) bind the N-lobe of CDK4 and CDK6 on the face opposite the cyclin, misaligning the catalytic residues and substantially reducing the kinase-cyclin interface (Brotherton et al., 1998; Jeffrey et al., 2000; Russo et al., 1998). CIP/KIP proteins (p21, p27 and p57) use a bipartite kinase-inhibitory domain in which a RxL segment engages the cyclin hydrophobic patch and a downstream 3_10_ helix inserts into the ATP site of CDK2 (Lacy et al., 2004; Russo et al., 1996; Wade Harper, 1993).

The CKI and CDK substrate interaction sets overlap almost completely outside the catalytic site. The cyclin hydrophobic patch was in fact defined by binding a motif shared by substrates and p21-like inhibitors and was thus considered a multipurpose docking site (Adams et al., 1996; Schulman et al., 1998). Similarly, yeast Sic1 engages Clb-Cdk through an RxL motif (Hodge and Mendenhall, 1999; Kõivomägi et al., 2011). The Cln1/2 LP pocket that recruits substrates such as Ste5 and Ste20 is occupied and blocked by Far1 (Bhaduri and Pryciak, 2011; Pope et al., 2014). The Cks1 phospho-threonine pocket likewise serves both inhibitors and substrates, reading priming sites in substrates and the pT173 and pT306 in Sic1 and Far1, respectively (Gartner et al., 1998; Moreno-Torres et al., 2017; Venta et al., 2020). Contacts outside that pocket also contribute: three separable Cks1 surfaces are required for p27 ubiquitination (Sitry et al., 2002), while no analogous contacts have been mapped for any yeast CKI.

In several proteins the distinction whether the protein functions as a CDK inhibitor or a substrate is determined by phosphorylation state. Phosphorylation of Sic1 at T173 is necessary to completely block Clb-Cdk1 activity to stop phosphorylation of the di-phosphodegrons (Moreno-Torres et al., 2017; Venta et al., 2020). In Far1, phosphorylation by the pheromone-activated MAP kinase Fus3 at T306 is required for cell cycle arrest and was proposed to strengthen binding to G1 cyclin-Cdk1 complexes, whereas without pheromone signalling Cdk1 phosphorylates the S87-S91 di-phosphodegron leading to Far1 ubiquitination by SCF^Cdc4^ (Blondel et al., 2000; Faustova et al., 2021; Gartner et al., 1998).

Far1 determines whether yeast cells divide or arrest in G1 to mate, acting as either an inhibitor or a substrate of the CDK complex. It inhibits Cln-Cdk1 *in vitro* (Peter and Herskowitz, 1994), and its only well-defined molecular function so far is blockade of cyclin-substrate docking at the Cln2 LP pocket (Pope et al., 2014). Far1 behaves is a substrate for S-phase CDK: its NLxxxL motif targets it to Clb5/6-Cdk1 for degron phosphorylation (Faustova et al., 2021). Clb5 is a key driver of S phase, yet overexpressing it does not let *cln1Δ cln2Δ cln3Δ* cells escape pheromone arrest (Oehlen et al., 1998). This indirectly suggests that Far1 may inhibit Clb5 in *vivo*. No structure of Far1 bound to a CDK complex exists, and neither an inhibition constant nor a kinetic mechanism has been reported for its inhibition of Clb5-Cdk1-Cks1. We therefore set out to identify the protein-protein interactions between Far1 and the CDK complex that are mediating its inhibitory function.

Here we report a previously undescribed interaction between the Far1 region ^310^QLIRTADISC^319^ and Cks1-Cdk1 that is required for CDK inhibition. We show that the isoleucines at positions 312 and 317 are important for Far1 mediated inhibition of the Clb5-Cdk1 complex, and that this region lowers the maximal velocity of the kinase while promoting multiphosphorylation Therefore, this Far1 region represents another possibility for regulation of protein multisite phosphorylation, with possible relevance in CDK substrates and inhibitors.

## Results

### Far1 region C-terminal of Cks1-bound pT306 contacts Cks1-Cdk1

We modelled the interactions between Clb5-Cdk1-Cks1 and Far1 with AlphaFold3, which suggested an interaction between Far1 ^310^QLIRTADISC^319^ region and Cks1-Cdk1 (**Figure 1A**). The identified ^310^QLIRTADISC^319^ region extends further than the known Cks1-interacting phospho-threonine motifs found in CDK substrate proteins (Kõivomägi et al., 2013; McGrath et al., 2013). We hypothesized that this interaction with Cks1-Cdk1 may be important for the Far1-mediated inhibition of CDK. To test that, we used recombinant Far1 fragments which we called Far1-min, because contain the Far1 region 280-393 that is sufficient region for CDK1 inhibition (Gartner et al., 1998; Reményi et al., 2005) (**Figure 1B**). As Far1 requires phosphorylation at T306 to induce its inhibitory activity (Far1 priming) (Gartner et al., 1998), we added a Fus3 docking motif (^72^KRGNIPKPLNL^82^ in Far1) to the N terminus of Far1-min and used analog-sensitive Fus3 to phosphorylate all Far1 variants in this work (**Figure 1C**).

**Figure 1.**
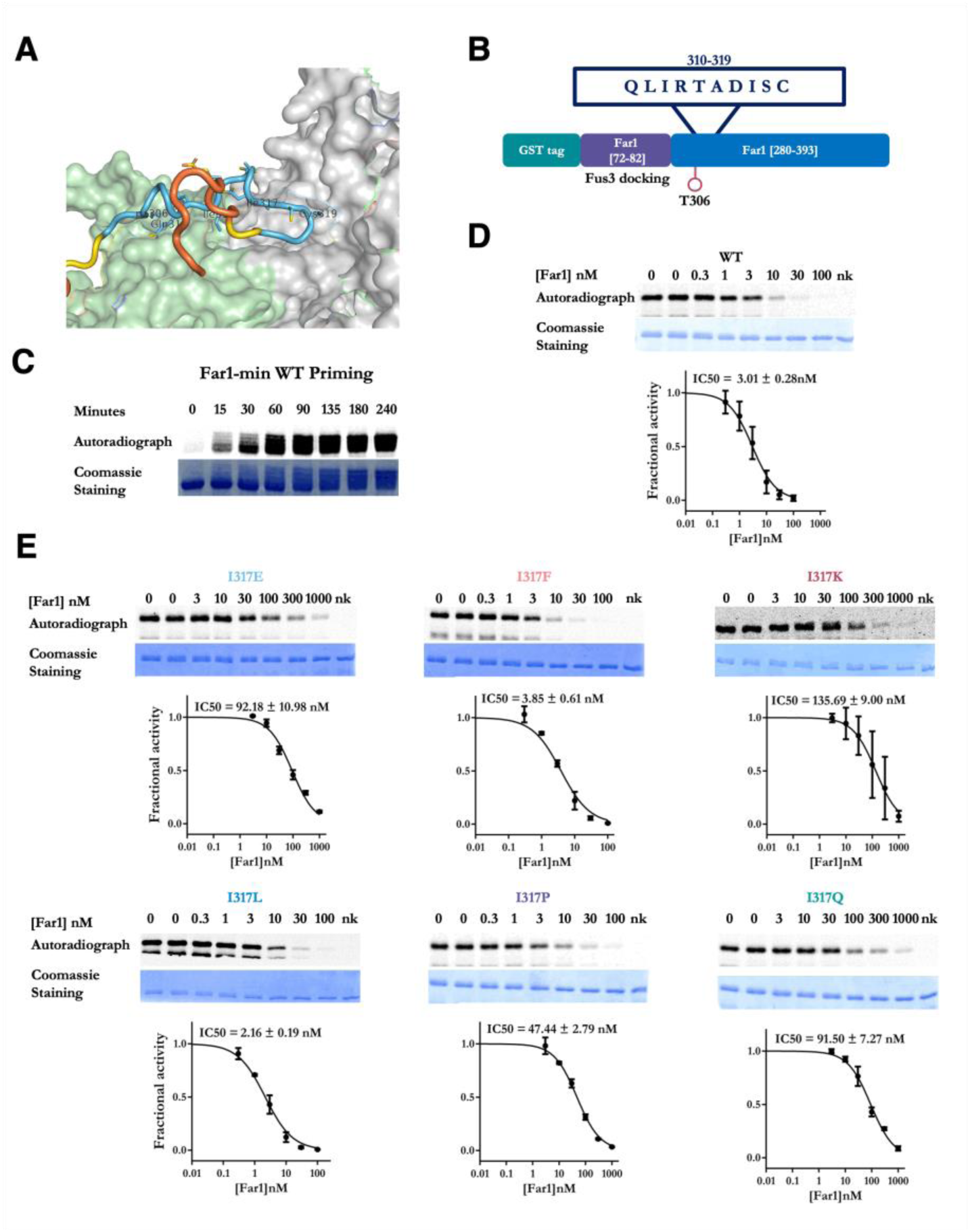
Far1 ^310^QLIRTADISC^319^ region interacts with Clb5–Cdk1-Cks1 and mutations in it disrupt CDK inhibition. **A.** Far1 residues Q310–C319 at the interface between Cks1 (green surface) and Cdk1 (grey surface) in the AlphaFold3 model of the Clb5–Cdk1–Cks1– pT306-Far1 complex. Far1 residues are coloured by per-residue model confidence (pLDDT): dark blue, very high (pLDDT > 90); light blue, confident (70–90); yellow, low (50–70); orange, very low (< 50). **B.** Schematic representation of the Far1-derived protein used in the assay, highlighting the Far1 ³¹⁰QLIRTADISC³¹⁹ linker sequence. **C.** ³³P autoradiograph and Coomassie-stained SDS-PAGE gel showing the time course of Far1 priming phosphorylation by Fus3. Samples were collected at 0, 15, 30, 60, 90, 135, 180, and 240 min after reaction initiation. **D, E.** Representative ³³P autoradiographs, Coomassie-stained SDS-PAGE gels, and corresponding IC_50_ plots for the WT Far1 construct (panel **‘D’**) and the different I317 mutants (panel **‘E’**). Assays were performed using a range of Far1 protein concentrations as inhibitor, Sic1ΔC as substrate and Clb5-Cdk1 as kinase. The 0 condition indicates reactions without Far1 and serves as a baseline control, whereas “nk” indicates reactions lacking the CDK complex (negative control). IC_50_ plots show data from at least two independent replicates, with error bars representing the standard error of the mean (SEM).

The Far1-Cks1-Clb5-Cdk1 AlphaFold model suggests that Far1 I317 is buried when bound to the CDK complex, and thus likely contributing to the interaction (**Figure 1A**). To test the model, we performed mutational scanning on I317 by mutating it to amino acids with different properties: negatively charged glutamic acid (E), hydrophobic amino acid with aromatic ring phenylalanine (F), positively charged amino acid lysine (K), hydrophobic amino acid leucine (L), conformationally rigid amino acid proline (P), polar uncharged amino acid glutamine (Q). We carried out *in vitro* kinase inhibition assays to determine how these point mutations affect the inhibition of CDK and measured half-maximal inhibitory concentration (IC_50_) for every mutant (**Figure 1 D, E**). We found that the wild-type version of Far1-min inhibited Clb5-Cdk1 with IC_50_ of 3 nM (**Figure 1D**). While mutating I317 to other hydrophobic residues (L, F) had little effect on IC50, mutation to all non-hydrophobic residues decreased the inhibitory potency considerably, with I317K mutation increasing IC50 45-fold (Figure 1E). This experiment shows that the region is relevant in Clb5-Cdk1 inhibition by Far1.

Next, we expanded the mutational scanning to the whole Far1 ^310^QLIRTADISC^319^ region and found that the substitutions in I312 (**Figure S4**) and I317 led to profound increase in Clb5-Cdk1 IC_50_ values, suggesting that these two amino acids are critical for the tight protein-protein interaction between Far1 and the CDK complex. I312 and I317 are also among the most deeply buried positions according to the AlphaFold3 model of the CDK-Far1 complex (**Figure 2B**). Out of 10 amino acid positions analysed, I312 and I317 were the only ones at which polar and hydrophobic substitutions behave differently: charged substitutions weaken inhibition whereas F and L are essentially comparable to the wild-type residue. Positions predicted to be solvent-exposed (R313, T314, A315, D316) show no strong preference for specific residues. Q310 and C319 are predicted to be buried but are mutationally tolerant; at Q310 the apparent effect derives from a single substitution (E, 12-fold; L 1.4-fold, Q 1.0-fold) (**Figure 2C**). Overall, our experimental findings agree with AlphaFold3 predicted model and demonstrate that this previously undescribed Far1 region plays a role in CDK inhibition.

**Figure 2.**
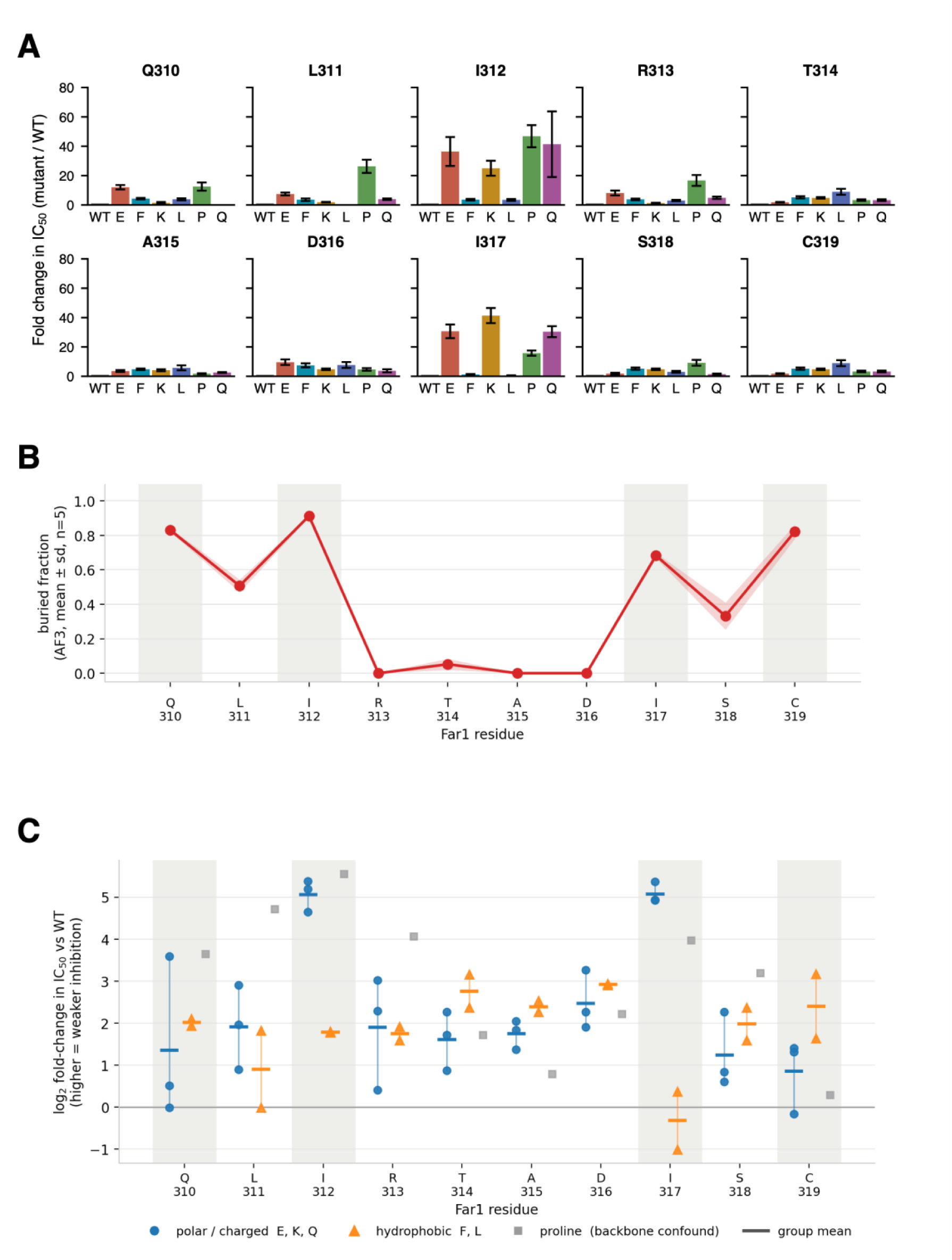
AlphaFold3 interface burial compared with the measured effect of Far1 substitutions on Cdk1 inhibition. **A.** Fold changes in IC_50_ values of the tested mutants relative to the WT Far1-derived protein, grouped by residue position. Bar charts show data from at least two independent replicates, with error bars representing the standard error of the mean (SEM). **B**. Buried fraction on complex formation for Far1 residues Q310–C319, calculated from the AlphaFold3 model of the Clb5–Cdk1–Cks1–pT306-Far1 complex (Shrake–Rupley ΔSASA; mean of five models, shaded band ± SD). Values near 1 indicate residues occluded by the Cdk1/Cks1 surface on binding; values near 0 indicate residues that remain solvent-exposed. Grey columns mark the positions with buried fraction > 0.6. **C.** Effect of single substitutions at the same positions on Far1 inhibition of Cdk1, expressed as log_2_ fold change in IC_50_ relative to wild-type Far1; higher values denote weaker inhibition. Each substitution is plotted by type: polar/charged (E, K, Q; blue), hydrophobic (F, L; orange) and proline (grey).

### Far1 QLIRTADISC region promotes multisite phosphorylation in Sic1 context

Next, we aimed to test whether the described ^310^QLIRTADISC^319^ Far1 region would affect Cdk1-mediated phosphorylation in a CDK substrate context. A previous study mapping the interactions involved in Cks1 phospho-docking in two different protein contexts found that the distance between priming and secondary site determines multisite phosphorylation efficiency, while the specific sequence of region between the Cks1 binding site and the secondary phosphorylation site plays limited role (Kõivomägi et al., 2013). Here, we tested if this specialized sequence from Far1 following the T306 Cks1 binding site could affect Cks1-dependent multiphosphorylation. For that we used a modified non-inhibitory Sic1 with only 2 phosphorylation sites (Sic1_2TP_ΔC) and introduced QLIRTADISC sequence between the first and second phospho-site (Sic1_2TP_ΔC_QLIRTADISC) (**Figure 3A**).

**Figure 3.**
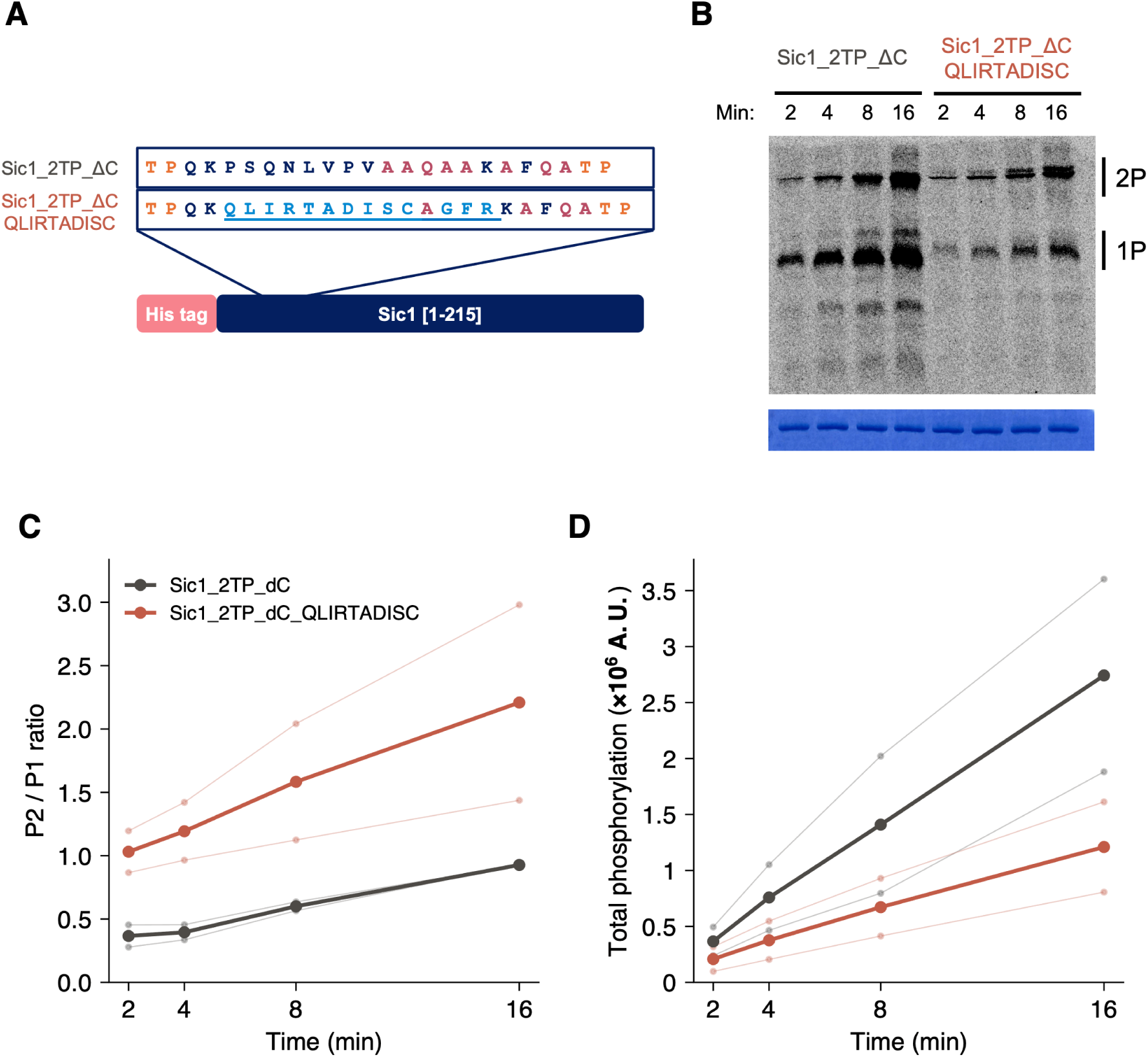
A Far1-derived sequence between Cks1 priming site and secondary CDK site alters the rate and processivity of phosphorylation. **A.** Schematic of the substrate used in the assay with highlighted Far1(QLIRTADISC) linker sequence between two CDK phosphorylation sites. **B.** Representative ^33^P autoradiograph of Phos-tag SDS-PAGE gels showing substrate phosphorylation by Clb5-Cdk1. Below, Coomassie staining as loading control showing the BSA band. **C.** Ratio of doubly and singly phosphorylated (P2/P1) products over four timepoints in *in vitro* kinase assays with Clb5-Cdk1. **D.** Total phosphorylation signal of both phosphoforms. Thinner lines show data from individual replicates, and thicker lines represent averages over replicates in panels **C** and **D**.

We carried out *in vitro* kinase assays using Cks1-Clb5-Cdk1 and found that QLIRTADISC region promotes multisite phosphorylation, leading to more than 2-fold increase in the doubly phosphorylated product, as measured by the mobility of the phospho-products in the Phos-tag SDS-PAGE (**Figure 3B, C**). We observed no multisite phosphorylation when using a phosphor-pocket mutant Cks1 (**Figure S11**). This result indicates that the interaction between this Far1 region and Cks1–Cdk1 is insufficient to compensate for the loss of phospho-pocket binding to the priming site, suggesting that the QLIRTADISC extends the phospho-threonine-containing Cks1 binding motif, but does not provide sufficient binding to function as an independent SLiM. This is also consistent with the previously reported essential role of T306 in Far1-mediated inhibition of the CDK complex (Gartner et al., 1998). Interestingly, despite the increasing in multiphosphorylation, the total phosphorylation rate was lower in Sic1_2TP_ΔC_QLIRTADISC (**Figure 3B, D**), hinting the inhibitory role of the linker region, as also seen in the Far1 context (**Figure 1, 2**).

### Far1 LIRTADISC region decreases substrate turnover rate in Sic1 context

To explore the mechanism behind the increased multisite phosphorylation but decreased total phosphorylation of the Sic1_2TP_ΔC_QLIRTADISC substrate by Clb5-Cdk1, we determined the Michaelis constant (K_m_) and maximal velocity (V_max_) values in a kinase assay (**Figure 4A, B**). Introduction of QLIRTADISC Far1 region to the Sic1 synthetic construct led to over two-fold decrease in V_max_ (**Figure 4C**) but did not affect the K_m_ of tested substrate (**Figure 4D**). Together with the increased multisite phosphorylation (**Figure 3**), this suggests that the QLIRTADISC-containing protein becomes more potent in engaging or retaining the kinase complex, but less effective at catalytic turnover. Alternatively, it is possible that a fraction of Clb5-Cdk1 complex is inhibited by Sic1_2TP_ΔC_QLIRTADISC, leaving a smaller fraction active to drive substrate phosphorylation. Together, this data indicates that providing additional contacts with the CDK complex between the canonical substrate recognition motifs changes the phosphorylation dynamics and may gradually shift a substrate protein towards a more inhibitor-like protein.

**Figure 4.**
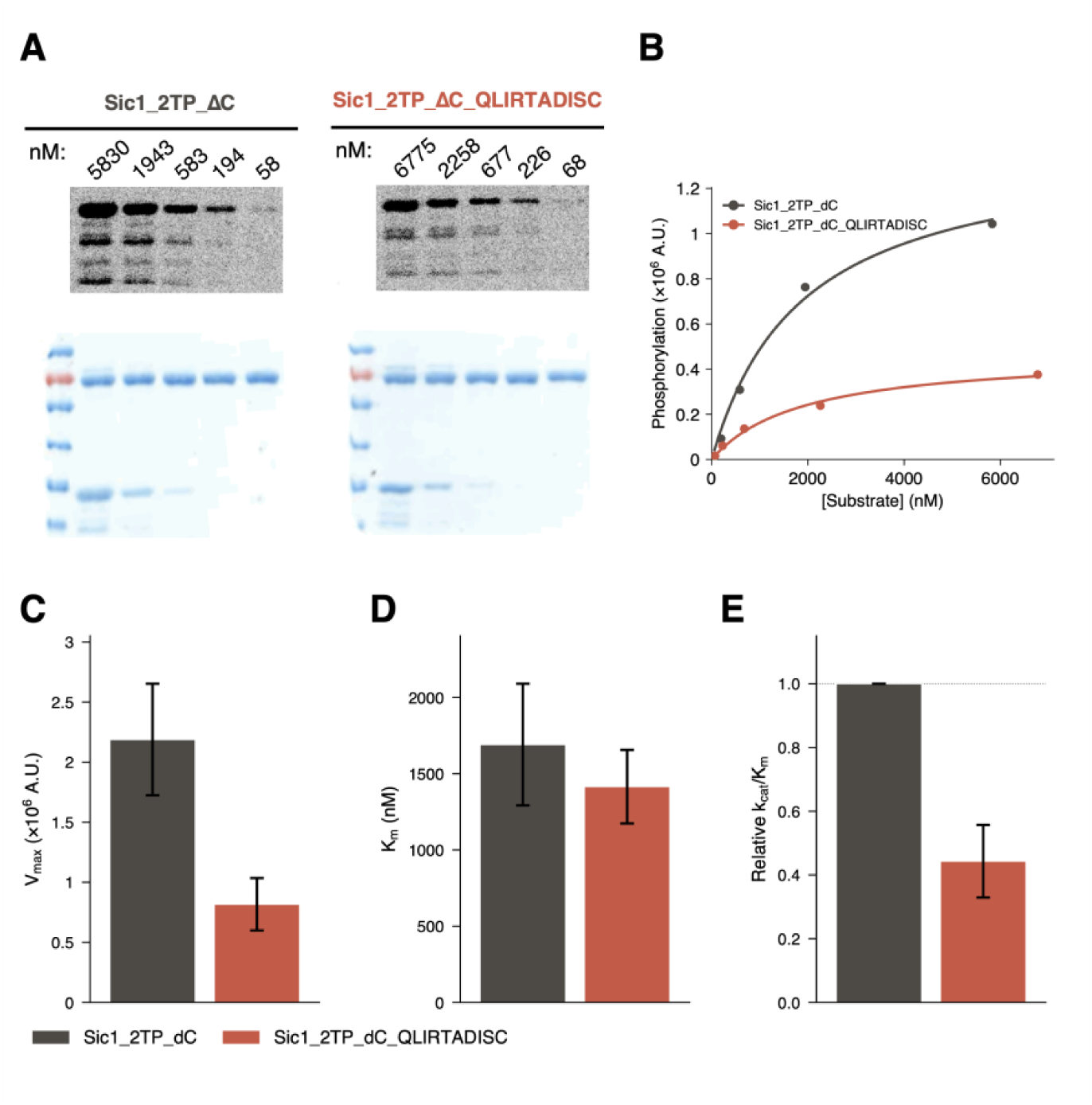
Far1-derived sequence between Cks1 priming site and secondary CDK site lowers phosphorylation rate through V_max_. **A.** Representative ^33^P autoradiograph of SDS-PAGE gels showing Sic1-based substrate phosphorylation by Clb5-Cdk1 *in vitro*. Below, Coomassie staining as loading control showing the BSA band (top) and the Sic1-based substrate (bottom). **B.** Representative Michaelis-Menten kinetics plot, Y-axis shows the phosphorylation rate (A.U.), X-axis is substrate protein concentration (nM). **C, D, E.** Bar charts showing V_max_, K_m_, and relative k_cat_/K_m_, respectively; mean of three replicates are shown with error bars indicating SEM.

## Discussion

Shared docking mechanisms contribute to both CDK substrate phosphorylation and inhibition, indicating that recognition of a docking motif alone does not determine the functional outcome of an interaction (Moreno-Torres et al., 2017; Örd et al., 2019; Venta et al., 2020). For Far1, the contacts that support inhibition beyond its established effects on cyclin-dependent substrate recruitment remain incompletely defined. Our results identify residues 310–319 ^310^QLIRTADISC^319^ as contributors to Clb5–Cdk1 inhibition, with I312 and I317 particularly important for inhibitory activity (**Figure 2A**). AlphaFold3 modelling predicts that these residues occupy shallow pockets near the Cks1–Cdk1 interface (**Figures 1A, 2B**), providing a possible structural explanation for their functional contribution. The model further predicts contacts between this Far1 region and Cks1 residues H29 and M45, as well as Cdk1 residues W176, S213 and D215. Although these predicted contacts remain to be experimentally validated, the agreement between the modelling and mutational data supports a role for this region in stabilising the Far1–CDK interaction. This extends the current understanding of Far1-mediated inhibition beyond its previously characterised interactions with cyclin docking surfaces.

The ^310^QLIRTADISC^319^ Far1 region seems to extend Cks1 phospho-threonine-centred SLiM and can promote multisite phosphorylation in a CDK substrate context (**Figure 3**) while slowing overall turnover. Since phospho-threonine-dependent docking for the multisite phosphorylation is still essential (**Figure S11**), it suggest that the Far1 310–319 region contributes an auxiliary interaction that acts together with the Cks1 phospho-pocket. Such cooperation could help explain how a short sequence contributes to inhibition in Far1 while altering phosphorylation-state distribution in a substrate context. The specific contacts and kinetic steps underlying these effects remain to be determined. Moreover, this auxiliary interaction of QLIRTADISC Far1 region may provide the mechanistic explanation for observed *in vivo* effect of partial restoration of inhibitory functionality of Far1-T306A, when it is stabilised by disrupting the Clb5-mediated phospho-degron. (Pope et al., 2014). Further, while selected mutations in the QLIRTADISC region caused up to 50-fold decrease in Far1-min inhibitory potential, introducing this region into Sic1 context did not turn Sic1 into a tight inhibitor, suggesting that Far1-min makes additional contacts to drive low nM inhibition of Clb5-Cdk1.

Our findings suggest that the distinction between CDK substrates and inhibitors may be influenced by interactions extending beyond their established docking motifs. Canonical cyclin-binding and Cks1-binding motifs provide important determinants of substrate recruitment and multisite phosphorylation, but the sequences connecting these elements can further tune the outcome of kinase engagement. Such auxiliary interactions could influence the balance between substrate phosphorylation, multisite phosphorylation and inhibition, providing an additional level of regulation for CDK-dependent signalling. Whether similar mechanisms operate in other yeast CDK inhibitors, or in human CDK regulatory proteins, remains an open question.

In conclusion, we identify a previously uncharacterised region of Far1 that contributes to CDK inhibition and modulates multisite phosphorylation when introduced into a substrate context. Our findings highlight the importance of sequences between canonical docking motifs and suggest that auxiliary interactions can influence both the extent of multisite phosphorylation and the overall rate of substrate processing. This provides an additional element in the molecular toolbox through which cyclin–CDK complexes regulate substrate recognition and phosphorylation.

## Acknowledgements

This work was funded by the Estonian Research Council grant PRG2149. This work received co-funding from the European Union’s Horizon Europe research and innovation programme under Grant Agreement No. 101060066 (DIGIBIO). It was also supported by the European Union through the European Regional Development Fund (Centre of Excellence for Bioengineering, project No. 2021-2027.1.01.24-0451) and by the Estonian Ministry of Education and Research. Views and opinions expressed are however those of the author(s) only and do not necessarily reflect those of the European Union or the European Research Executive Agency (REA). Neither the European Union nor the granting authority can be held responsible for them.

The Authors would like to thank Regina Virre for her excellent technical assistance and dedicated support in maintaining the laboratory infrastructure throughout this study. The authors would like to thank Mart Loog for making this project possible.

The authors used ChatGPT (OpenAI) for text editing, grammatical polishing of the whole document, Claude Science Opus 5 (Anthropic) for literature discovery support and polishing the Introduction of the manuscript. The authors reviewed and edited all outputs and take full responsibility for the final content of the manuscript.

## Author contributions

**AM:** Conceptualization, Data curation, Formal analysis, Investigation, Project administration, Software, Supervision, Validation, Visualization, Writing – original draft

**JM:** Data curation, Formal analysis, Investigation, Validation, Visualization

**VK:** Data curation, Investigation

**EV:** Conceptualization, Methodology, Supervision

**ML:** Formal analysis, Visualization, Writing – original draft

**MÖ:** Conceptualization, Supervision, Writing – review & editing

**IF:** Conceptualization, Funding acquisition, Methodology, Project administration, Resources, Supervision, Writing – review & editing.

## Methods

### Cloning

All Far1 variants were cloned into pGEX4T-1 vector using Inverse PCR cloning method described in (Silva et al., 2017). All plasmids used are listed in supplementary Table S1.

### Protein purification

Clb5-Cdk1 was purified from yeast cells using TAP method with C-terminally tagged cyclins as described previously (Puig et al., 2001; Ubersax et al., 2003). Cks1 was purified according to (Reynard et al., 2000). GST-tagged Far1 variants and analog-sensitive version of Fus3(Q93G) (Bishop et al., 2000) were expressed in *E. coli* BL21RP cells at 16°C using 0.5 mM IPTG. For purification of GST-tagged proteins, Glutathione Sepharose (GE Healthcare) was used. GST-tagged analog-sensitive version of Fus3(Q93G) (Fus3-as) was co-purified with Ste5 peptide (Bhattacharyya et al., 2006). His-tagged Sic1ΔC (1-215) was expressed in *E. coli* BL21RP at 37°C using 2 mM IPTG and purified by using Chelating Sepharose (GE Healthcare) loaded with CoCl2 and eluted with 200 mM imidazole.

### Priming of Far1 variants with Fus3-as

After purification, all Far1 variants were phosphorylated with Fus3-as for activation (below referred to as Far1 priming). The composition of the priming mixture contained 5 mM MgCl_2_, 2 mM ATP, 150 nM Fus3-as, and 6 μM Far1 variant. The reaction was carried out at room temperature for 4 h and stopped by adding 1-(1,1-dimethylethyl)-3-(1-naphthalenyl)-1H-pyrazolo[3,4-d] pyrimidin-4-amine (1-NA-PP1) to the final concentration of 40 μM. To follow the phosphorylation process, 40 μL of the reaction mixture was transferred to a separate tube, the radioactive ATP ([γ-^33^P]-ATP) (Hartmann Analytic) was added to the mix, and timepoints at 0, 15, 30, 60, 90, 135, 180, 240 minutes after reaction initiation were collected to assess Far1 phosphorylation. The samples were run on SDS-PAGE, the gels were stained in Coomassie R-250 staining solution (0,1 % Coomassie Blue R-250 (w/w), 30 % methanol, 5 % acetic acid), destained in destaining solution (30 % methanol, 5 % acetic acid) and dried. Amersham Typhoon 5 biomolecular Imager (GE Healthcare Life Sciences) was used to detect radioactive signals from storage phosphor screens. The quantification of signals was performed in ImageQuant TL (Amersham Biosciences). Microsoft Excel and GraphPad Prism version 10.2.0 were used for data analysis. Primed Far1 variants were desalted before the *in vitro* kinase inhibition assays using 7K MWCO ZebaTM Spin Desalting Columns (Prod# 89882) according to the instructions provided by the manufacturer.

### *In vitro* kinase inhibition assays

The general composition of the kinase inhibition assays was the following: 50 mM Tris-HCl (pH 7.4), 150 mM NaCl, 5 mM MgCl_2_, 20 mM imidazole, 2% glycerol, bovine serum albumin (BSA) (0.2 mg/ml), 500 nM Cks1, and 500 μM adenosine triphosphate (ATP) [(with added [γ-^33^P]-ATP (Hartmann Analytic)], 10 μM 1-NA-PP1. Sic1ΔC (1-215) at the final concentration of 1 μM was used as a substrate in all experiments. To evaluate IC50 values, we carried out 7 reactions with Far1 variant concentrations ranging from 0.3 nM to 1.2 μM, as well as two reactions without Far1 to obtain the baseline phosphorylation rate. The concentration of the Clb5-Cdk1 complex was 0.2 nM. Before the initiation of reaction, Clb5-Cdk1 complex was incubated with primed Far1 variant for 10 minutes at room temperature and then reaction was started by adding Sic1ΔC (1-215) substrate. After 16 minutes at room temperature, reactions were stopped by adding SDS-PAGE sample buffer to reaction mixture. The samples were run on SDS-PAGE, the gels were stained in Coomassie R-250 staining solution (0,1 % Coomassie Blue R-250 (w/w), 30 % methanol, 5 % acetic acid), destained in destaining solution (30 % methanol, 5 % acetic acid) and dried. Amersham Typhoon 5 biomolecular Imager (GE Healthcare Life Sciences) was used to detect radioactive signals from storage phosphor screens. The quantification of signals was performed in ImageQuant TL (Amersham Biosciences). Microsoft Excel and GraphPad Prism version 10.2.0 were used for data analysis. All *in vitro* kinase inhibition assays were performed at least twice.

### *In vitro* kinase assays

The general composition of the kinase assays was the following: 50 mM Tris-HCl (pH 7.4), 150 mM NaCl, 5 mM MgCl_2_, 20 mM imidazole, 2% glycerol, bovine serum albumin (BSA) (0.2 mg/ml), 500 nM Cks1, and 500 μM adenosine triphosphate (ATP) [(with added [γ-^33^P]-ATP (Hartmann Analytic)]. For substrate, proteins based on the non-inhibitory fragment of Sic1 were used at the final concentration of 1 μM. The concentration of the Clb5-Cdk1 complex was 0.2 nM. The phosphorylation reaction was initiated by adding the kinase to the substrate. Samples were collected after 8 and 16 minutes (in some cases 2, 4, 32 minutes) by mixing the 8 μL of the reaction mixture with SDS-PAGE sample buffer. For electrophoresis, the samples were then loaded to 10% SDS-PAGE gels, or 8% Phos-tag SDS-PAGE gels with 100 mM Phos-tag and 200 mM MnCl2. The Phos-tag SDS-PAGE gels were prepared as described in (Örd and Loog, 2020). Phos-tag gels were used to determine the multisite phosphorylation. Amersham Typhoon 5 biomolecular Imager (GE Healthcare Life Sciences) was used to detect radioactive signals from storage phosphor screens. The quantification of signals was performed in ImageQuant TL (Amersham Biosciences). Microsoft Excel and GraphPad Prism version 11.0.2 were used for data analysis. All *in vitro* kinase assays were performed at least twice.

### Relative K_cat_/K_m_ calculations

The kinase assays were performed as described above. Substrates were serially diluted in approximately half-log_10_ steps (alternating 3.0- and 3.33-fold dilutions, i.e. a 10-fold decrease every two steps) across eight concentrations. Samples were collected at 8 and 16 minutes after reaction initiation. Phosphorylation was linear in time between 8 and 16 minutes (signal_16_/signal_8_ ≈ 2) across the substrate concentration range, confirming initial rate conditions for the kinetic analysis.

The phosphorylation signal was plotted against substrate concentration and fitted to the Michaelis–Menten equation by non-linear regression (GraphPad Prism version 11.0.2), fitting each biological replicate independently; 95% confidence intervals were estimated by the profile-likelihood method.

Because the highest attainable substrate concentration only modestly exceeded Km, Vmax and Km were individually weakly constrained; catalytic efficiency (k_cat_/K_m_, proportional to V_max_/K_m_) was therefore used as the robust comparator. k_cat_/K_m_ are reported relative to the control substrate (Sic1_2TP_dC = 1). The control was assayed in every experiment, and each variant was normalized to the control measured in the same experiment, cancelling between-experiment differences in signal scale (e.g. exposure time). Relative kcat/Km is reported as the mean ± SEM of n = 3 independent experiments.

### Structure prediction

The quaternary complex of Clb5 (UniProt P30283, residues 1–435), Cdk1 (P00546, 1–298), Cks1 (P20486, 1–150) and full-length Far1 (P21268, 1–830) was predicted with AlphaFold3 via the AlphaFold Server (alphafoldserver.com). T306 of Far1 was specified as phosphothreonine. The job returned five predicted models that were used for analysis. Structures were rendered in PyMOL 3.1.0.

### Buried-surface analysis

Per-residue solvent-accessible surface area (SASA) was computed with the Shrake–Rupley algorithm as implemented in Biopython 1.84 (Bio.PDB.SASA.ShrakeRupley, probe radius 1.4 Å, 100 sphere points). For each residue of Far1, SASA was calculated twice: once for the isolated Far1 chain and once for Far1 within the full Clb5–Cdk1–Cks1 complex. The buried fraction on complex formation was defined as: buried fraction = 1 – SASA (in complex) / SASA (isolated chain). Values were computed independently for each of the five models and are reported as mean ± standard deviation.

### Data Visualization and Software

Data visualizations (Figures 2A, 3C, 3D, 4B, 4C, 4D, 4E) were generated using custom Python scripts. An artificial intelligence tool (Claude 4.8 Opus, Anthropic) was utilized to assist in writing, refining, and debugging the code scripts. All data processing and figure rendering were executed locally on a private workstation; no experimental datasets were uploaded to or processed by external AI servers.

